# A pilot study of the comparison using a standard and individual altitude exposure protocol in elite gymnasts

**DOI:** 10.1101/2023.12.08.569719

**Authors:** A.A. Grushin, I.E. Zelenkova, D. Ilin, S.V. Zotkin, P.V. Korneev, S.V. Koprov, D.Kh. Almyashev, V.A. Badtieva, O.S Glazachev

## Abstract

**Aim:** Evaluate the effectiveness of the use of altitude training in elite gymnasts and compare two approaches of the altitude exposure protocols.

**Materials and methods:** The study involved 6 athletes of high qualification specializing in gymnastics (22 (20.5-22.5 years, 166.5 (162.5-169) cm, 63.5 (60-65 kg)). Athletes underwent hypoxic exposure using “Live high - train low” method using standard (18 days) and individual protocol (18 days). The biochemical parameters of blood, total hemoglobin mass, blood volume, hypoxic index, and oxygen saturation of arterial blood were determined.

**Results:** The tHb-mass values before the standard protocol were 678.5 (664-712.5) g, immediately after 696 (679-752.7) g. Before the start of the individual protocol, the tHb-mass values were 687 (622.7-752) g, immediately after 751.5 (750.2-827) g, (p < 0.05). The tHb-mass shifts after exposure to the unified protocol were 24 (4.8-31) g, and after personified 73.5 (55.5-132.7) g, (p < 0.05). At the same time, ALT, AST, and CPK values did not differ from the baseline, and testosterone values increased after a personalized hypoxic exposure program.

**Conclusions:** The application of personalized protocol of additional hypoxic impact seems to be more effective than standard protocol in terms of change of hematological parameters.

## Introduction

Scientific studies on training in natural altitude with the aim to improve performance can be attributed to the 60s of the last century. This was primarily related due to the celebration of the international competitions at altitude: the Winter Olympic Games at an altitude of 1900 meters in Squaw Valley in 1960 and the Summer Olympic Games in Mexico City in 1968 at an altitude of 2200-2300 meters above sea level. With the accumulation of scientific evidence training at altitude began to be actively used not only to increase performance at altitude but also at sea level. Increase in performance is related not only to cardiorespiratory and haematological mechanisms of adaptation, such as a change in hemodynamic characteristics, ventilatory function of the lungs and others ^1^, but also due to peripheral (tissue and intracellular) adaptation mechanisms such as increased muscle buffer capacity, accelerated resynthesis and creatine phosphate reserves, increased glycolytic power ^2^, cost-effectiveness, and others. ^3^

Understanding the basic mechanisms of short-term and long-term adaptation to hypoxia made it possible to develop methods for using hypoxic training not only in sports with the predominant contribution of aerobic systems but also with a high anaerobic contribution, for example, team sports and others. Gymnastics is considered a complex coordination sport in which a change in all components of coordination abilities can be associated with fatigue. In this regard, maintaining high performance (fatigue resistance) is a specific component of endurance that is directly related to the gymnastics technique. In a study by Sawczyn S et al., it was shown that gymnasts have high aerobic performance associated with a decreased pulse during the performance of intense gymnastic combinations and with slowing the development of “coordination fatigue.” ^4^

This can be of particular importance at the elite level with a large amount of training load and many new complex combinations are mastered. Numerous studies show that one of the most effective methods of hypoxic exposure is the approach on the principle of “Live high -train low” and “Live high - train lower and higher.” ^56 7^. Traditionally, in the training of elite athletes, a unified approach is used, in which athletes are exposed to the same altitude. Despite the fact that this approach is considered quite practical-oriented, a personification of hypoxic effects is advisable. This is due to the fact that the reaction to hypoxic effects varies significantly from individual to individual, depending on the current athlete’s physiological state, period of preparation, etc. ^8^. For example, in a study by Friedmann et al. (2005) it was shown that the increase in erythropoietin concentration after 4 hours in normobaric hypoxia (FiO_2_=15%, approximately 2500 m) was on average ∼ 58%, but noticeably ranged from 10 to 185%. ^9^. In the study of Nummela et al., regression analysis showed that exposure height was the most significant factor explaining the positive increase in total haemoglobin mass. In addition, it was shown that male athletes had a higher total haemoglobin mass gain than female athletes. ^10^.

For the same athletes, training at altitude can be effective, or maybe not, which can be related to altitude. The introduction of individualization in hypoxic exposure is actively discussed in the scientific community and various approaches have already been proposed to assess the necessary and effective hypoxic dose. ^11 12 13 14 15^

In 2012, within the framework of a special project of the ROC Innovation Center, a personalized approach to the application of additional hypoxic effects was developed and tested, based on the “Live high - train low” technique for athletes specializing in endurance sports which was used in the study ^16^.

The main purpose of this study was to evaluate the effectiveness of the use of altitude training in elite gymnasts in order to increase aerobic capacity and compare the standard and individual protocol based on the approach «Live high - train low».

## Materials and Methods

Six elite gymnasts (22 (20.5-22.5) years, 166.5 (162.5-169) cm, 63.5 (60-65) kg) members of the National team participated in the study. Study design - balanced longitudinal cross-sectional. First, all 6 athletes were doing normobaric hypoxic training (using hypoxic tents and hypoxic generators) according to the standard protocol for 18 days (ST), and 60 days after the end of the exposure, they underwent the program of hypoxic exposures modified according to individualisation algorithm (IND). The 60-day wash-out period was related to the time course of the total haemoglobin mass stabilisation after the altitude exposure with the goal of elimination the effect of the first experiments on the results of the second experiment.^17^ All the athletes underwent a similar training program according to the preparation phase and trained at least 24 h.wk−1.

During the first 18 days of ST hypoxic exposure started with an altitude of 1500 m above sea level (FIO_2_ = 17%) and daily altitude was increased by 300 m (respectively, reducing the concentration of oxygen in the inhaled air) to reach an altitude of 3000 m above sea level. Further, the altitude was maintained at 3000 m above sea level (FIO_2_ = 14%) (Table 1).

**Table 1.**
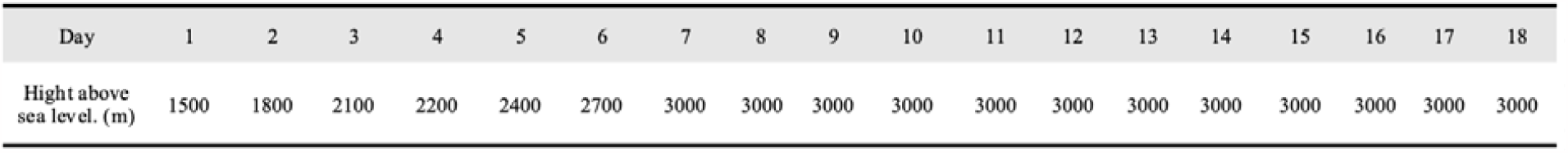
Protocol of unified program of additional hypoxic exposure (n = 6, 18 days). maintained at 3000 m above sea level (FIO2 = 14%).

The normobaric hypoxia exposure using the approach “Live high - train low” was carried out using the Hypoxico Everest Summit II generator (Hypoxico, USA), designed to obtain hypoxic gas mixtures by membrane separation of ambient air and hypoxic tents, where the athlete was in the evening, during night sleep and daytime rest. On average, the daily hypoxic exposure was 10-11 hours. In the morning, immediately after waking up inside the tent athletes daily recorded blood saturation values with oxygen saturation (SpO_2_) and HR using a pulse oximeter MD300C318 (Beijing Choice Electronic Tech Co., Ltd, Beijing, PRC).

IND hypoxic exposures were carried out according to the original algorithm based on the values of the following physiological parameters: SpO_2_, the recovery index, and the results of a hypoxic test using a ballroom system. SpO_2_ was measured in the morning same as in the ST approach.

The recovery index was taken from the night measurement using the Bodyguard device (Firstbeat, Finland) to register overnight R-R intervals, the data was analyzed using the Firstbeat SPORTS computer program provided by the manufacture. Based on the native recording, the software calculates the recovery index (RI). The recovery index is based on the analysis of the heart rate variability (HF and LF). It is reported as a percentage of reduction from 100% ^18^

The hypoxic test was carried out using a Hypoxico Everest Summit II generator (Hypoxico, USA), and pulsoximeter MD300C318 (Beijing Choice Electronic Tech Co., Ltd, Beijing, PRC). Briefly, athletes in the sitting position were breathing normobaric hypoxic air (6400 m, FIO2 10.1±0.1%) till SpO2, drops down to 80%. After arterial oxygen saturation rich 80% or doesn’t rich 80% during 300-sec subject switched to breathing with atmospheric air till SpO2-rich 96%. For the calculation of the hypoxic index time of reaching 80% is divided by the time of restitution to 96% of SpO2. ^19^.

All three parameters were evaluated in a complex using a table and a ballroom system (minimum score - 0, maximum - 6). The reference values for the ballroom system: I-Hyp: 0-1.9 c.u. - 0 points, 2-3 c.u. - 1 point, >3 c.u. - 2 points; RI 0-39 % - 0 points, 40 - 69 %— 1 point, 70-100% 2 points; SpO2 >87% - 0 points, 88-93 %— 1 point, <94% - 2 points. ^16^. If the athlete has 0 points in the sum of three indicators, then the height should be “lowered” by 300 m above s.l.; if one point 200 m above the s.l.; 2-3 points - the athlete during the next hypoxic session “remains” at the same height; 4 points - you should “raise” the height of the hypoxic exposure by 100 m above s.l.; 5 - 200 m above s.l.; 6 - 300 m above s.l. ^16^. If an athlete is feeling sick or changes in the biochemical blood analysis, the altitude was adjusted according to the results of the consultation and joint expert decision by a team doctor, coach, and sports physiologist.

Before and right after the ST and IND exposure following tests were performed: biochemical tests, total hemoglobin mass (tHb-mass) and blood volume (BV) determination, and hypoxic test (Figure 1). Additionally, several biochemical markers were measured: testosterone, cortisol, urea, CPK, ALT, and AST were used to determine the individual response to training loads and hypoxic exposures.

**Figure 1.**
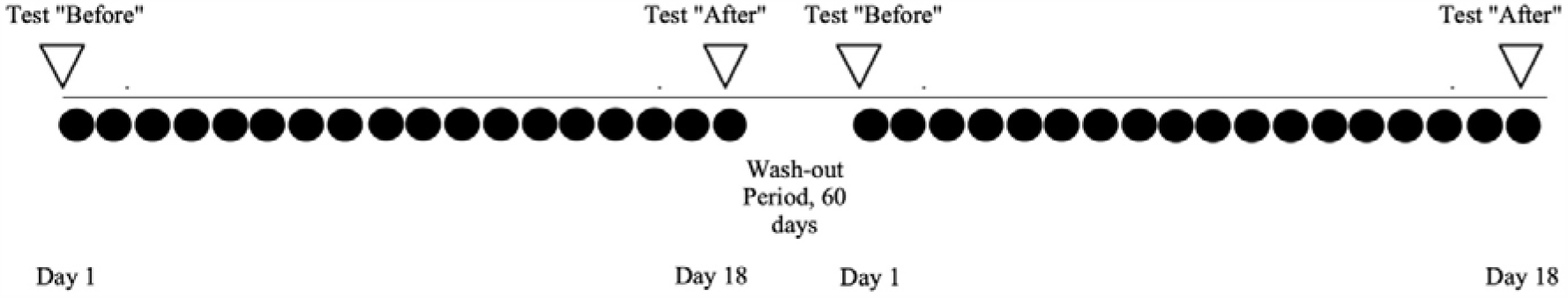
Study design.

The measurement of tHb-mass and BV was carried out using the carbon monoxide (CO) breath-back technique ^20^. For 2 minutes, the athlete breathed through a gas glass spirometer (Bloodtec, Germany) with a mixture containing CO (1 ml/kg for M). To determine hemoglobin CO saturation (% HbCO), capillary blood (210 μL) was taken before the procedure, at the 6th and 8th minutes of return breathing. Blood samples were analyzed by CO-oximeter (ABL80 FLEX COOX Radiometer, Denmark). The calculation of amount of CO before and 4 minutes after the start of the procedure was measured for the amount of SO_2_ in exhaled air using an analyzer (Drager Pac 800, Germany). For 2 minutes, the athlete breathed through a spirometer of a gas glass spirometer (Bloodtec, Germany) with a mixture containing CO (1 ml/kg for M, 0.8 ml/kg for G). To determine hemoglobin CO saturation (% HbCO), capillary blood (210 μL) was taken before the procedure, at the 6th and 8th minutes of return breathing. Blood samples were analyzed by CO-oximeter (ABL80 FLEX COOX Radiometer, Denmark). The calculation of the tHb-mass and circulating BV was carried out using software provided by the developer of this technique (Bloodtec, Germany) ^20^. heart rate and heart rate variability (HF and LF). It is reported as a percentage of maximum reduction ^21^. Hemoglobin concentration and hematocrit were determined by Hemo Control (EKF, Germany). Capillary blood was used for the analysis. Capillary blood was used to determine the serum concentration of cortisol, urea, CPK, ALT, and AST (DRG kit, Germany). To quantify cortisol in human blood serum by enzyme immunoassay on a solid-phase carrier, a spectrophotometer (BTS-350, Spain) eppendorphic centrifuge, dosers (10 μl to 5 ml), and a microplate shaker were used. The microtiter plate reader was measured for 15 minutes. The reference data, according to the manufacturer, was a concentration of 160 to 770 nmol/l. To determine the serum testosterone concentration a set of reagents (DRG, Germany) to quantify testosterone in human blood serum by enzyme immunoassay on a solid-phase carrier, and spectrophotometer (BTS-350, Spain) were used. Briefly, the desired number of coated wells to be used was placed in the holder. 25 μl of each standard, control, and sample were pipetted into the respective wells. 100 μl of conjugate reagent was added to each well. All samples were excavated for 15 minutes. Thoroughly mixed for 30 seconds. Incubated for 60 minutes at 18-23ºC. The wells were washed with wash solution 3 times and their contents were shaken. 100 μl of TMV reagent was added to each well. The mixture was gently stirred for 10 seconds and incubated for 20 minutes at room temperature (18-25 ° C). The reaction was stopped by adding 100 μL of the solution to each well. The microtiter plate reader was measured for 15 minutes. Reference data, according to the manufacturer, were the concentration from 9.0 to 42 nmol/l.

Statistical data processing was carried out using Prism 9 from GraphPad. Since the sample size was small and the distribution differed from normal, median and interquartile spans were used to characterize the samples. The paired Wilcoxon test was used to compare the bound samples. The significance level was 0.05.

Informed consent was obtained from all subjects. All methods were carried out in accordance with relevant guidelines and regulations. All experimental protocols were approved by the First Medical University Ethic Commission.

## Results

The application of ST hypoxic exposure resulted in significant changes in the values of various haematological and biochemical parameters (Table 2).

**Table 2.** Summary table of measured parameters during standard (ST) and individualised protocol (IND) (n = 6).

|  | ST |  | IND |  |
| --- | --- | --- | --- | --- |
|  | Before | After | Before | After |
| **Hemoglobin, g/l** | 149 (146-152) | 151 (150-162) | 162 (149-163) | 162 (151,7-164) |
| **Hematocrit, %** | 44 (44-45) | **45 (44-48) #** | 48 (47-48) | 48 (45-49) |
| **Serum iron, mcmol/L** | 15,4 (14,7-15,7) | 19,2 (13,9-29,5) | 27,6 (26,7-31,9) | 19,5 (18,7-29,2) |
| **Total hemoglobin mass, g** | 678,5 (664-712,5) | 696 (679-752,7) | 687 (622,7-752) | **751,5 (750,2-827) \*** |
| **Testosterone/cortisol, s.u.** | 6,2 (5,7-6,8) | 5,1 (3,9-6,0) | 5,1 (3,8-5,7) | **6,2 (5,5-7,2) \*** |
| **Cortisol, nmol/l** | 265 (211-299) | **453 (433-502) \*** | **575 (535-684) #** | **591,5 (574,8-604) \*** |
| **Testosterone, nmol/l** | 15,2 (14,4-16,6) | 21 (18,4-28,2) | **26,1 (24,9-26,6) #** | **33 (28,3-37,9) \*#** |
| **Creatine phosphokinase, E/L** | 246 (156-270) | 189 (180-293) | 371 (308-575) | 240 (194,5-254) |
| **Alanine transferase, E/L** | 21 (17-22) | 18 (14-22) | 21 (17-22) | 17 (16-18) |
| **Aspartateaminotransferase, E/L** | 39 (37-39) | **31 (25-33) \*** | 33 (31-33) | 30,5 (25,7-33) |

\*- difference from baseline, $p < 0.05$

ST approach for all athletes did not lead to a reliable increase in tHb-mass but was accompanied by a significant increase in cortisol values and a tendency to increase testosterone values.

The use of hypoxic exposure using the IND approach led to a reliable increase in tHb-mass. At the same time, Htc values by the end of the second stage were significantly higher than in the same period of the first stage of the study. The values of cortisol and testosterone concentration, before IND significantly increased compared with the data before the ST.

Individual tHb-mass values before and after hypoxic exposure programs are presented in Figure 2 and Figure 3. The average tHb-mass gain after ST was 24 (4.8-31) g, and after IND was 73.5 (55.5-132.7) g, (p < 0.05). Due to the fact that the personalized protocol is unique for each athlete and varies from individual to individual, Figures 4 provide examples of individual hypoxic exposure programs. Recovery index values were recorded only during the application of the personalized hypoxic exposure protocol and were 69 (66-83)% on the first day after program initiation and 53 (48-73)% and did not differ. The average RI change values in the average group were −16 (−18- (−9))%.

**Figure 2.**
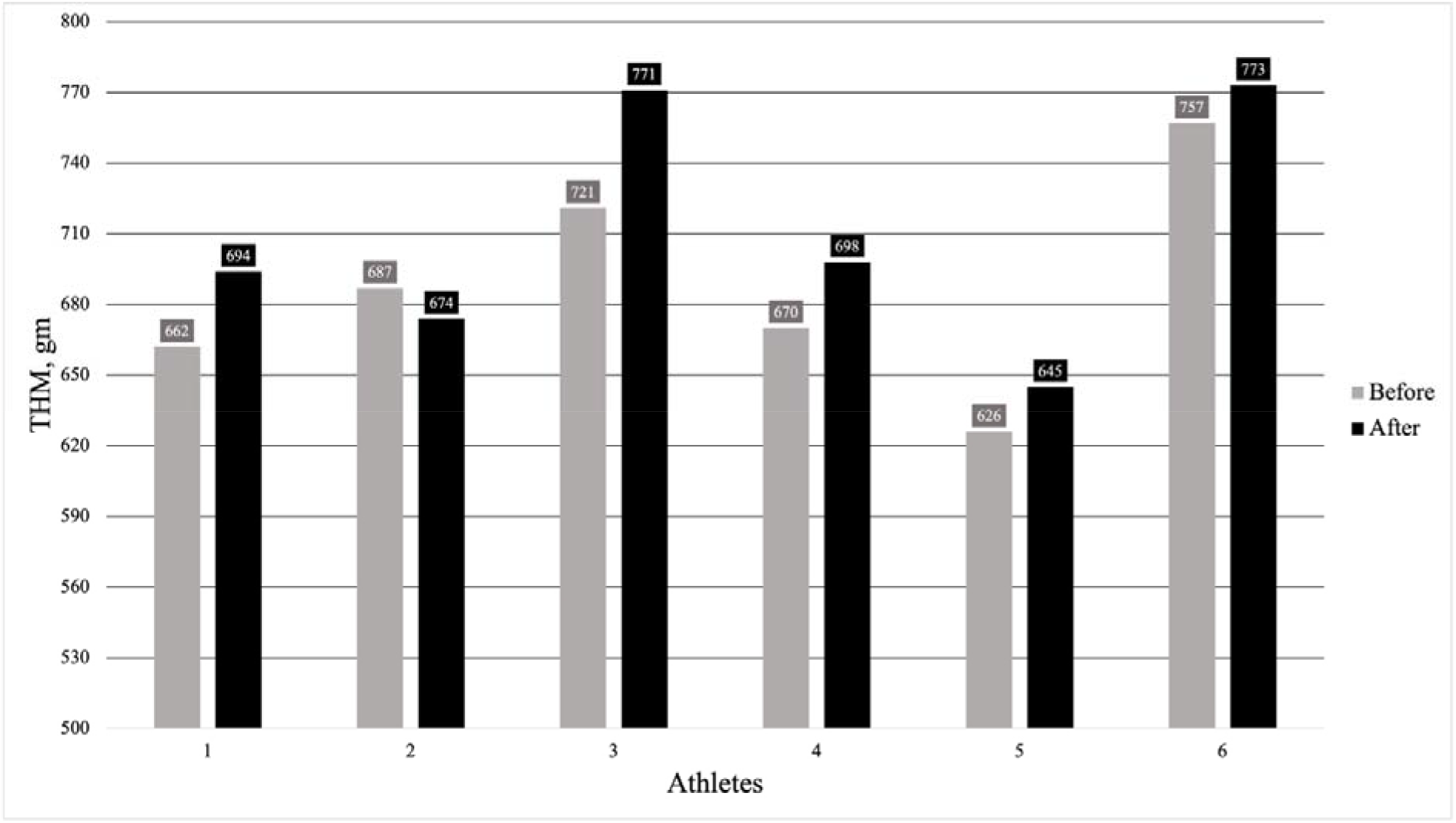
Individual tHb-mass values before and after the ST.

**Figure 3.**
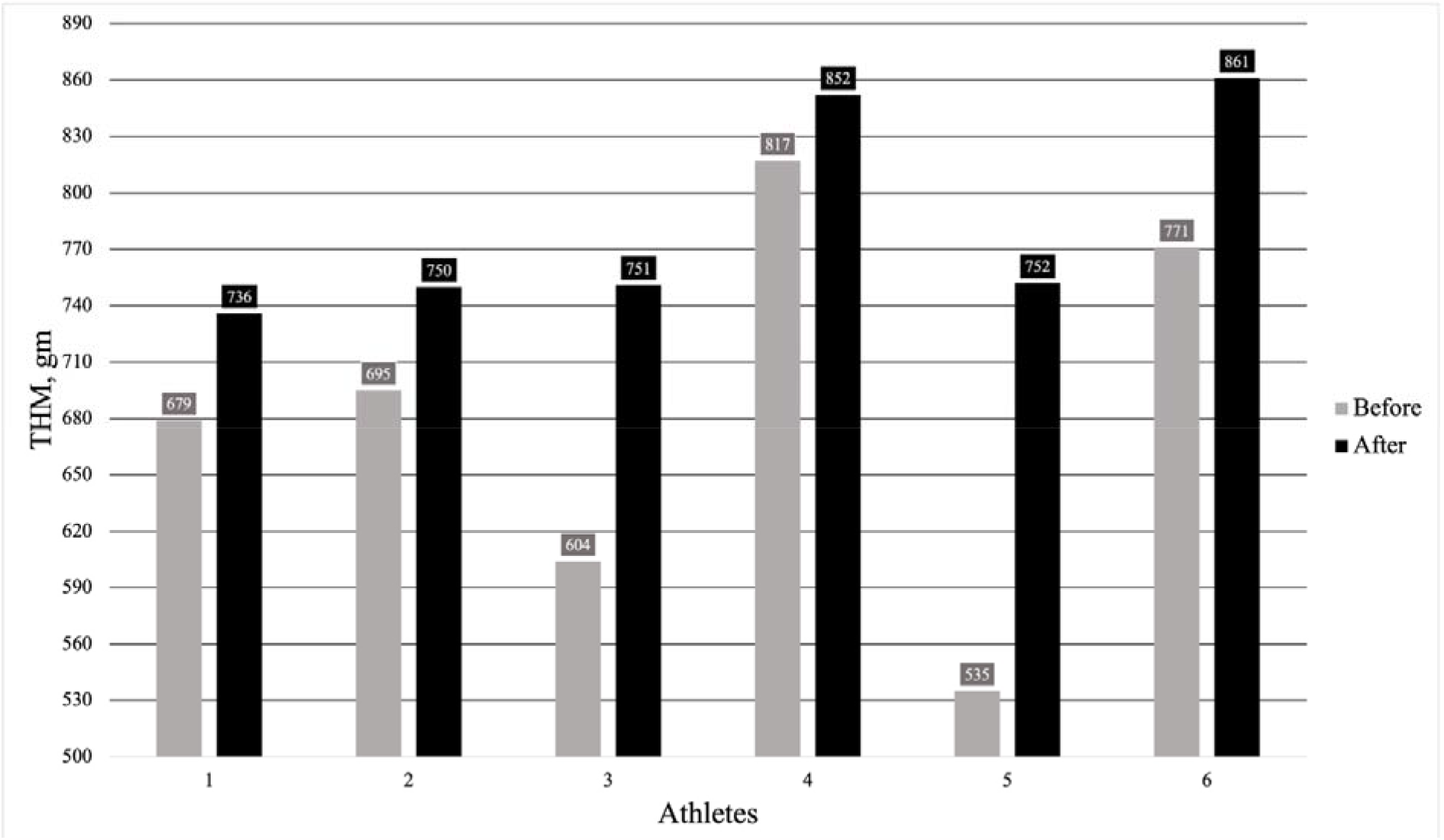
Individual values of tHb-mass before and after IND.

**Figure 4.**
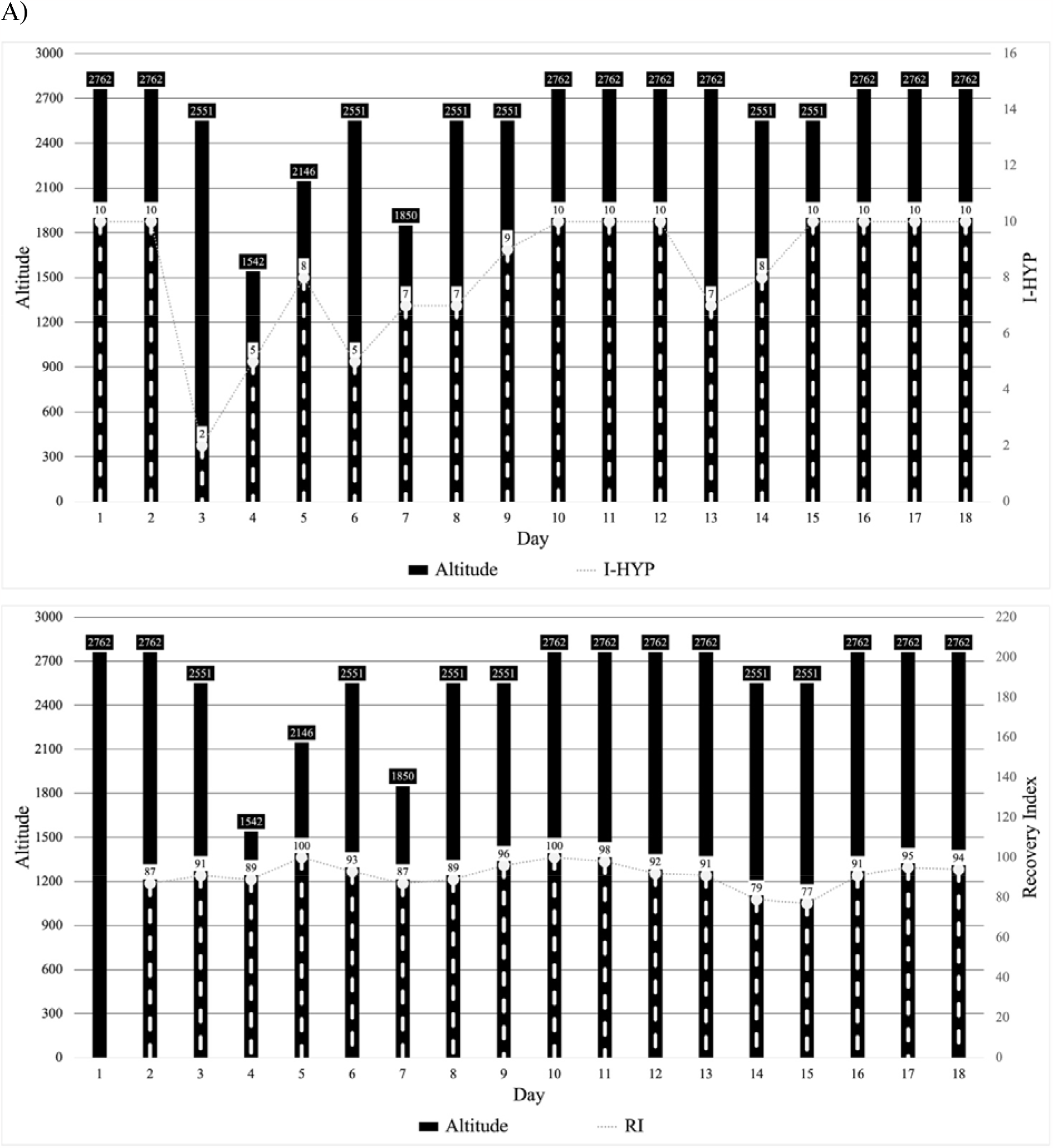
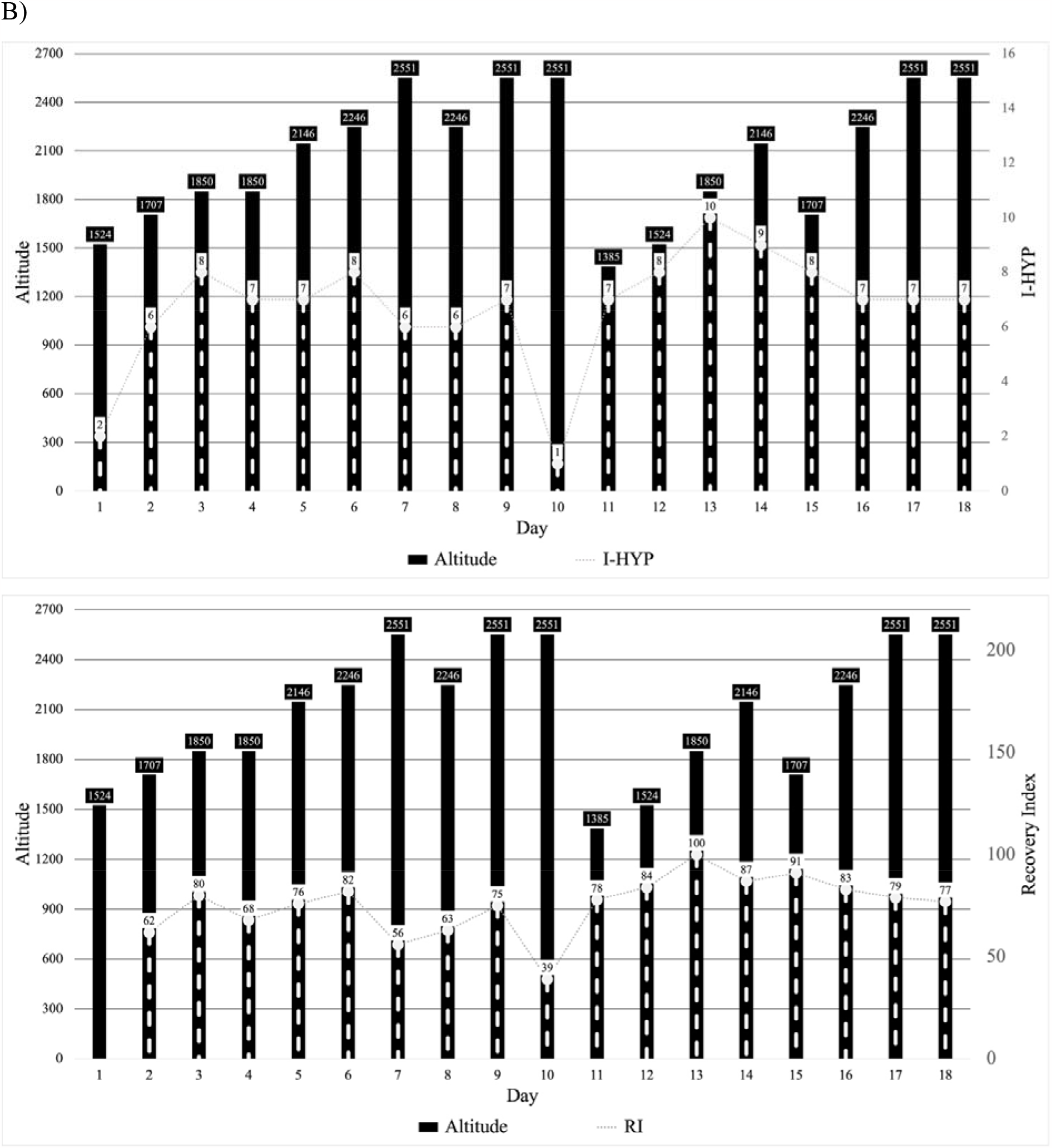
Protocols of the individual program: athlete 1 (A) and athlete 2 (B).

The average cortisol gain after the ST was 203 (68-204) nmol/l, and after IND - 89 (60-52) nmol/l (p < 0.05). The average cortisol increase was 6.6 (5.5-11.6) nmol/L after ST and 12.6 (11.8-17.3) nmol/L after IND and did not differ between the groups. The average values of the degree of increase in the ratio of testosterone to cortisol in the group after ST were −1.8 (−1.8 - (- 0.1) s.u. and 5.3 (1.6 - (−2.9) s.u., (p < 0.05) after IND.

## Discussion

To our knowledge, this is the first study related to the application of hypoxic exposure in gymnastics. In the context of gymnastics, it is further worth emphasizing that reduced aerobic performance can lead to “coordination fatigue,” and additional interventions to increase aerobic performance in gymnasts reduce HR during basic exercise according to listed research ^4^. The application of hypoxic exposure in gymnastics can be effective in order to increase performance and reduce the risk of developing “coordination fatigue.” Unfortunately in our study, we were not able to register HR during the training but self reported fatigue was lower in after hypoxic exposure.

In order to compare the influence of the different hypoxic exposure protocols on aerobic performance tHb-mass values were used. According to studies, there is a high correlation between tHb-mass and VO2 max, with no such relationship for Hb and Hct. ^22^. tHb-mass has also been shown to be more stable compared to Hb and Hct. For example, the concentration of hemoglobin is highly dependent on the BV, which in turn can lead to incorrect results and does not allow measuring absolute values ^23 24^. Hct level is also a very labile indicator and depends on the change in blood plasma volume ^25^. The absence in the present study of a significant increase in tHb-mass may be associated with insufficient duration of hypoxic exposure. However, comparing tHb-mass changes in two experiments the increase after the IND protocol was observed, with can suggest in favor of the personalised approach.

In order to assess how the body of the athletes adapt to hypoxic exposure, biochemical blood parameters were analyzed. The ratio of testosterone to cortisol in fasting morning is often used in the literature to characterize the ratio of anabolic to catabolic processes in the body ^26^. This attitude reflects, first of all, the body’s adaptation response to the load presented in the previous few days. It is shown that after heavy training loads, this ratio will decrease, and after lighter loads, it can even increase slightly. According to previous studies, the response of cortisol to exercise has been shown to depend on both the duration and intensity of the load ^27^. It has been suggested that the critical threshold above which circulating cortisol levels rise corresponds to an exercise duration of more than 20 minutes at about 60-70% maximum oxygen intake ^26 27^. Gymnastics training is characterized by periodic “bursts” of physical activity interspersed with rest and recovery periods. Given that gymnasts use most of their energy through anaerobic pathways with little contribution from aerobic ^28^, this fact may serve as an explanation for the increase in cortisol concentration in both groups ^29 30^. After applying a personalized hypoxic exposure, cortisol values were higher than after a unified protocol. On the one hand, this may be related to the use of hypoxic exposure and the predominance of catabolic processes. On the other hand, baseline of the cortisol before the IND was higher than before ST which also can explain the difference after the exposure. Additionally from the results of the study, we see that testosterone values in the group that applied the individual approach increased, and other biochemical markers do not differ between groups. Thus, the increase in cortisol level can be associated with the training cycle period that took place between two hypoxic exposure periods. According to R.M. Daly á et al., gymnasts undergo a significant increase in cortisol during the transition from the base to the pre-competition cycle, which corresponds to the training plan: a unified hypoxic exposure program was used in the base cycle, and personified in the pre-competition. ^31^ In this case, the cortisol values in our study are similar to those of the study mentioned above. An increase in other biochemical markers such as ALT, AST, and CPK weren’t recorded. This suggests that the use of additional hypoxic exposure during intensive training activities does not lead to increased cell damage in both muscle and liver tissues.

According to the results of the study, it was shown the use of the personified protocol of hypoxic exposure in gymnastics is more effective than the unified protocol in terms of changing hematological parameters, and also does not cause a pronounced increase in catabolic processes during intensive training activities.

## References

1. Sinex JA, Chapman RF. Hypoxic training methods for improving endurance exercise performance. J Sport Health Sci. 2015;4(4):325–332. doi:10.1016/j.jshs.2015.07.005

2. Sahlin K. Muscle energetics during explosive activities and potential effects of nutrition and training. Sports Med. 2014;44 Suppl 2:S167–73. doi:10.1007/s40279-014-0256-9

3. Jung W-S, Kim S-W, Park H-Y. Interval Hypoxic Training Enhances Athletic Performance and Does Not Adversely Affect Immune Function in Middle- and Long-Distance Runners. Int J Environ Res Public Health. 2020;17(6). doi:10.3390/ijerph17061934

4. Sawczyn S, Biskup L, Zasada M, Mishchenko V. Special endurance of young gymnasts: the role of aerobic capacity in fatigue development in the training. Central European Journal of Sport Sciences and Medicine. 2018;23:59–70. doi:10.18276/cej.2018.3-06

5. Sharma AP, Saunders PU, Garvican-Lewis LA, et al. Training Quantification and Periodization during Live High Train High at 2100 M in Elite Runners: An Observational Cohort Case Study. Journal of sports science & medicine. 2018;17(4):607–616.

6. Levine BD, Stray-Gundersen J. “Living high-training low”: effect of moderate-altitude acclimatization with low-altitude training on performance. J Appl Physiol. 1997;83(1):102–112. doi:10.1152/jappl.1997.83.1.102

7. Brocherie F, Millet GP, Hauser A, et al. “Live High-Train Low and High” Hypoxic Training Improves Team-Sport Performance. Med Sci Sports Exerc. 2015;47(10):2140–2149. doi:10.1249/MSS.0000000000000630

8. Chapman RF, Stager JM, Tanner DA, Stray-Gundersen J, Levine BD. Impairment of 3000-m run time at altitude is influenced by arterial oxyhemoglobin saturation. Med Sci Sports Exerc. 2011;43(9):1649–1656. doi:10.1249/MSS.0b013e318211bf45

9. Friedmann B, Frese F, Menold E, Kauper F, Jost J, Bärtsch P. Individual variation in the erythropoietic response to altitude training in elite junior swimmers. Br J Sports Med. 2005;39(3):148–153. doi:10.1136/bjsm.2003.011387

10. Nummela A, Eronen T, Koponen A, Tikkanen H, Peltonen JE. Variability in hemoglobin mass response to altitude training camps. Scand J Med Sci Sports. 2021;31(1):44–51. doi:10.1111/sms.13804

11. Soo J, Girard O, Ihsan M, Fairchild T. The use of the spo2 to fio2 ratio to individualize the hypoxic dose in sport science, exercise, and health settings. Front Physiol. 2020;11:570472. doi:10.3389/fphys.2020.570472

12. Vinogradov M, Zelenkova I. Dose-response modelling of total haemoglobin mass to hypoxic dose in elite speed skaters. BioRxiv. Published online June 18, 2020. doi:10.1101/2020.06.18.159269

13. Pla R, Brocherie F, Le Garrec S, Richalet J-P. Effectiveness of the hypoxic exercise test to predict altitude illness and performance at moderate altitude in high-level swimmers. Physiol Rep. 2020;8(8):e14390. doi:10.14814/phy2.14390

14. Gore CJ, Sharpe K, Garvican-Lewis LA, et al. Altitude training and haemoglobin mass from the optimised carbon monoxide rebreathing method determined by a meta-analysis. Br J Sports Med. 2013;47 Suppl 1:i31–9. doi:10.1136/bjsports-2013-092840

15. Bonetti DL, Hopkins WG. Sea-level exercise performance following adaptation to hypoxia: a meta-analysis. Sports Med. 2009;39(2):107–127. doi:10.2165/00007256-200939020-00002

16. Volkov AA. Methodical ROC recommendations. Application of additional artificial hypoxicimpact in sports of the highest achievements. Published online 2013:11.

17. Wachsmuth NB, Völzke C, Prommer N, et al. The effects of classic altitude training on hemoglobin mass in swimmers. Eur J Appl Physiol. 2013;113(5):1199–1211. doi:10.1007/s00421-012-2536-0

18. Firstbeat Technologies Ltd. Recovery Analysis for Athletic Training Based on Heart Rate Variability. In: Firstbeat Technologies Ltd.; 2015.

19. Zelenkova I, Zotkin S, Korneev P, et al. Individual variation in hypoxic tolerance across athletes of different level and sport specialisations. Gazz Med Ital - Arch Sci Med. 2020;179(3). doi:10.23736/S0393-3660.19.04045-2

20. Schmidt W, Prommer N. The optimised CO-rebreathing method: a new tool to determine total haemoglobin mass routinely. Eur J Appl Physiol. 2005;95(5-6):486–495. doi:10.1007/s00421-005-0050-3

21. Vesterinen V, Häkkinen K, Laine T, Hynynen E, Mikkola J, Nummela A. Predictors of individual adaptation to high-volume or high-intensity endurance training in recreational endurance runners. Scand J Med Sci Sports. 2016;26(8):885–893. doi:10.1111/sms.12530

22. Schmidt W, Prommer N. Impact of alterations in total hemoglobin mass on VO 2max. Exerc Sport Sci Rev. 2010;38(2):68–75. doi:10.1097/JES.0b013e3181d4957a

23. Schumacher YO, Pottgiesser T, Ahlgrim C, Ruthardt S, Dickhuth HH, Roecker K. Haemoglobin mass in cyclists during stage racing. Int J Sports Med. 2008;29(5):372–378. doi:10.1055/s-2007-965335

24. Convertino VA. Blood volume: its adaptation to endurance training. Blood volume: its adaptation to endurance trainingMed Sci Sports Exerc. 1991;23(12):1338–1348.

25. Schmidt W, Biermann B, Winchenbach P, Lison S, Böning D. How valid is the determination of hematocrit values to detect blood manipulations? Int J Sports Med. 2000;21(2):133–138. doi:10.1055/s-2000-8871

26. Urhausen A, Gabriel H, Kindermann W. Blood hormones as markers of training stress and overtraining. Sports Med. 1995;20(4):251–276. doi:10.2165/00007256-199520040-00004

27. Viru A. The pituitary-adrenocortical system. In: Hormones in muscular activity. CRC Press, Boca Ration, Florida. 1985;1(Hormonal ensemble in exercise):25–60.

28. Kirkendall DT. Physiologic aspects of gymnastics. Clin Sports Med. 1985;4(1):17–22.

29. Buono MJ, Yeager JE, Hodgdon JA. Plasma adrenocorticotropin and cortisol responses to brief high-intensity exercise in humans. J Appl Physiol. 1986;61(4):1337–1339. doi:10.1152/jappl.1986.61.4.1337

30. Elias AN, Wilson AF, Pandian MR, et al. Corticotropin releasing hormone and gonadotropin secretion in physically active males after acute exercise. Eur J Appl Physiol Occup Physiol. 1991;62(3):171–174. doi:10.1007/BF00643737

31. Daly RM, Rich PA, Klein R. Hormonal responses to physical training in high-level peripubertal male gymnasts. Eur J Appl Physiol Occup Physiol. 1998;79(1):74–81. doi:10.1007/s004210050476

